# Conserved and context-dependent roles for Pdgfrb signaling during zebrafish vascular mural cell development

**DOI:** 10.1101/2021.03.29.437552

**Authors:** Koji Ando, Yu-Huan Shih, Lwaki Ebarasi, Ann Grosse, Daneal Portman, Ayano Chiba, Kenny Mattonet, Claudia Gerri, Didier Y.R. Stainier, Naoki Mochizuki, Shigetomo Fukuhara, Christer Betsholtz, Nathan D. Lawson

## Abstract

Platelet derived growth factor beta and its receptor, Pdgfrb, play essential roles in the development of vascular mural cells, including pericytes and vascular smooth muscle. To determine if this role was conserved in zebrafish, we analyzed *pdgfb* and *pdgfrb* mutant lines. Similar to mouse, *pdgfb* and *pdgfrb* mutant zebrafish lack brain pericytes and exhibit anatomically selective loss of vascular smooth muscle coverage. Despite these defects, *pdgfrb* mutant zebrafish did not otherwise exhibit circulatory defects at larval stages. However, beginning at juvenile stages, we observed severe cranial hemorrhage and vessel dilation associated with loss of pericytes and vascular smooth muscle cells in *pdgfrb* mutants. Similar to mouse, *pdgfrb* mutant zebrafish also displayed structural defects in the glomerulus, but normal development of hepatic stellate cells. We also noted defective mural cell investment on coronary vessels with concomitant defects in their development. Together, our studies support a conserved requirement for Pdgfrb signaling in mural cells. In addition, these mutants provide an important model for definitive investigation of mural cells during early embryonic stages without confounding secondary effects from circulatory defects.

**Summary statement:** Genetic analysis in zebrafish demonstrates the conserved role of Pdgfb/Pdgfrb signaling in pericyte and vascular smooth muscle cell formation during vascular development in vertebrates.

## INTRODUCTION

During vasculogenesis in vertebrate embryos, endothelial cells initially form a primitive vascular network that subsequently becomes invested by mural cells (MCs) emerging *de novo* from the surrounding mesenchyme (Ando et al., 2019; Beck and D’Amore, 1997; Hungerford et al., 1996). Subsequently, pre-existing MCs co-migrate and co-proliferate with endothelial cells during angiogenesis to cover newly established blood vessels (Benjamin et al., 1998; Hellstrom et al., 1999). By morphology and gene expression, MCs are categorized into at least two cell types: pericytes and vascular smooth muscle cells (VSMCs) (Armulik et al., 2011; Vanlandewijck et al., 2018). Recent single-cell RNA sequencing of the mouse brain vasculature has revealed that pericytes and venous VSMCs form a phenotypic continuum distinguished by a progressive increase in the expression of contractile proteins in the venous VSMCs (Vanlandewijck et al., 2018). Arterial and arteriolar VSMCs, on the other hand, form a distinct continuum of gene expression patterns. Thus, the adult mouse brain vasculature appears to contain two classes of MCs, arterial/arteriolar VSMCs and pericyte/venous VSMCs, respectively, occupying distinct zones along the arterio-venous axis.

Similar to endothelial cells, pericytes specialize according to organ residence (Augustin and Koh, 2017; Muhl et al., 2020; Vanlandewijck et al., 2018). For example, mesangial cells reside within the kidney glomerulus where they contact endothelial cells and basement membrane to bridge glomerular capillary loops (Farquhar and Palade, 1962; Latta et al., 1960; Sakai and Kriz, 1987). Mesangial cells are considered specialized pericytes, but also have properties of SMCs (Schlondorff, 1987) and fibroblasts (He et al., 2021). In the liver, hepatic stellate cells, reside within the perisinusoidal space between the hepatocytes and sinusoidal endothelial cells and play a role in the storage of vitamin A (Yin et al., 2013), while in the brain, pericytes are essential for the function of the blood-brain barrier (Armulik et al., 2010). In each case, pericytes play an important role in defining and maintaining the organotypic function of the particular capillary bed with which they associate.

Analysis of mice lacking platelet-derived growth factor-B *(Pdgfb)* and its receptor, Pdgfrb, demonstrated their requirement for MC development (Hellstrom et al., 1999; Levéen et al., 1994; Lindahl et al., 1997; Soriano, 1994). Endothelial cells secrete Pdgfb, which activates Pdgfrb on neighboring MCs (Armulik et al., 2005). While initial MC specification is largely independent of Pdgfb or Pdgfrb, they are required for MC proliferation and recruitment to new vessels (Armulik et al., 2005; Hellstrom et al., 1999). *Pdgfb* and *Pdgfrb* mutant mice have also revealed insights into the requirements of MCs for endothelial development and vascular stabilization. *Pdgfb-* and *Pdgfrb*-deficient mice develop multiple vascular abnormalities, including structural defects in the glomerulus, dilation of heart and blood vessels, and extensive hemorrhage in numerous organs. Interestingly, VSMC coverage appears normal on major arteries in the absence of Pdgfb signaling despite defects in vascular stability. Due to the numerous functional defects, *Pdgfb* and *Pdgfrb* null mutants are perinatally lethal, preventing analysis of postnatal processes. Development of hypomorphic and conditional alleles, such as the *Pdgfb^ret/ret^, Pdgfb^EC-flox^* (Armulik et al., 2010; Lindblom et al., 2003), or *Pdgfrb*^F7/F7^ (Tallquist et al., 2003), have provided insights into the functional role of pericytes at postnatal stages (Armulik et al., 2010; Daneman et al., 2010; Vanlandewijck et al., 2015). However, a more detailed analysis of the effects of Pdgfb/Pdgfrb deficiency during early stages of embryonic development is challenging in mouse.

The zebrafish embryo is ideal for investigating cardiovascular development during embryogenesis. Zebrafish embryos are transparent and exhibit rapid external development, providing an accessible platform for visualizing the vascular system. Importantly, zebrafish embryos are small enough to limit secondary effects of hypoxia due to circulatory defects. These benefits allow more direct analysis of cellular and molecular defects when genetic manipulations lead to loss of circulatory function. Previously, we have generated fluorescent protein reporter lines driven by the *pdgfrb* and *abcc9* loci to directly visualize MCs in developing zebrafish (Ando et al., 2016; Ando et al., 2019; Vanhollebeke et al., 2015; Vanlandewijck et al., 2018). We have leveraged these lines to analyze and visualize the developmental dynamics of MCs during their recruitment to blood vessels. In the course of these studies, we presented a preliminary analysis of MCs in zebrafish embryos bearing a point mutation in *pdgfrb* (Ando et al., 2016). However, a more comprehensive phenotypic analysis is lacking. Here, using additional Pdgfb/Pdgfrb signaling mutants, together with multiple MC reporter lines, we investigate the role of Pdgfb signaling for MC recruitment and vascular maintenance in zebrafish. Our results demonstrate a conserved requirement for Pdgfb signaling in MC development, while providing a model to assess proximal cellular and molecular effects of Pdgfb signaling deficiency during embryonic development.

## RESULTS

### Pdgfb/Pdgfrb signaling is required for zebrafish brain pericyte development

We previously generated a mutation in *pdgfrb (pdgfrb^um148^)* that leads to nonsense-mediated decay of *pdgfrb* transcript and loss of Pdgfrb protein (Kok et al., 2015). Embryos mutant for *pdgfrb^um148^* or the ENU-induced null mutant, *pdgfrb^sa16389^*, appear morphologically normal until 5 dpf, with no obvious defects in circulatory function (Ando et al., 2016; Kok et al., 2015). To characterize possible MC defects in this mutant, we crossed *pdgfrb^um148^* onto the *TgBAC(pdgfrb:egfp)^ncv22^* background and assessed brain MCs at 5 dpf, with a focus on pericytes. As shown previously, *pdgfrb:egfp* is expressed in MCs covering central arteries throughout the brain at 5 dpf (**Fig. S1A, B**). Most *pdgfrb:egfp-positive* MCs on brain capillaries do not express smooth muscle markers, such as *Tg(acta2:mcherry)^ca8^* (**Fig. S1A, B**) or *TgBAC(tagln:egfp)* (Ando et al., 2016), consistent with their identity as pericytes. By contrast, *acta2:mcherry-positive* VSMC appear deeper in the brain vasculature at the Circle of Willis (CoW), a network of larger caliber vessels that integrate circulatory flow from the paired carotid arteries (**Fig. S1C**). These cells also express *pdgfrb:egfp*, although more weakly than pericytes (**Fig. S1A-C**), consistent with observations in mouse (Vanlandewijck et al., 2018). At 5 dpf, *pdgrb^um148^* larvae show a severe loss of *pdgfrb:egfp-positive* pericytes from the central arteries in the mid- and hindbrain when compared to homozygous wild type or heterozygous siblings (**Fig. 1A-C**). An exception was the metencephalic artery, which demarcates the boundary between the mid- and hindbrain (Isogai et al., 2001; arrowheads in **Fig. 1A, B**). We also noted a small decrease in cranial vessel volume in *pdgfrb^um148^* mutants (**Fig. 1D**), although mutants still showed severe pericyte loss when normalized to vessel volume (**Fig. 1E**). Observed pericyte loss could be due to failure of *pdgfrb:egfp* expression. Therefore, we assessed pericyte coverage of cranial arteries using transmission electron microscopy (TEM). In homozygous wild type siblings, we found that a majority of endothelial cells exhibited direct pericyte contact at their abluminal side and a shared basement membrane (**Fig. 1F, H**). In heterozygous *pdgfrb^um148^* siblings we noted a small decrease in endothelial cells with pericyte coverage (**Fig. 1H**). We failed to detect any endothelial cells in association with pericytes in *pdgfrb^um148^* mutants (**Fig. 1G, H**). Together, these results demonstrate that *pdgfrb^um148^* mutant embryos display a loss of brain pericytes.

**Figure 1.**
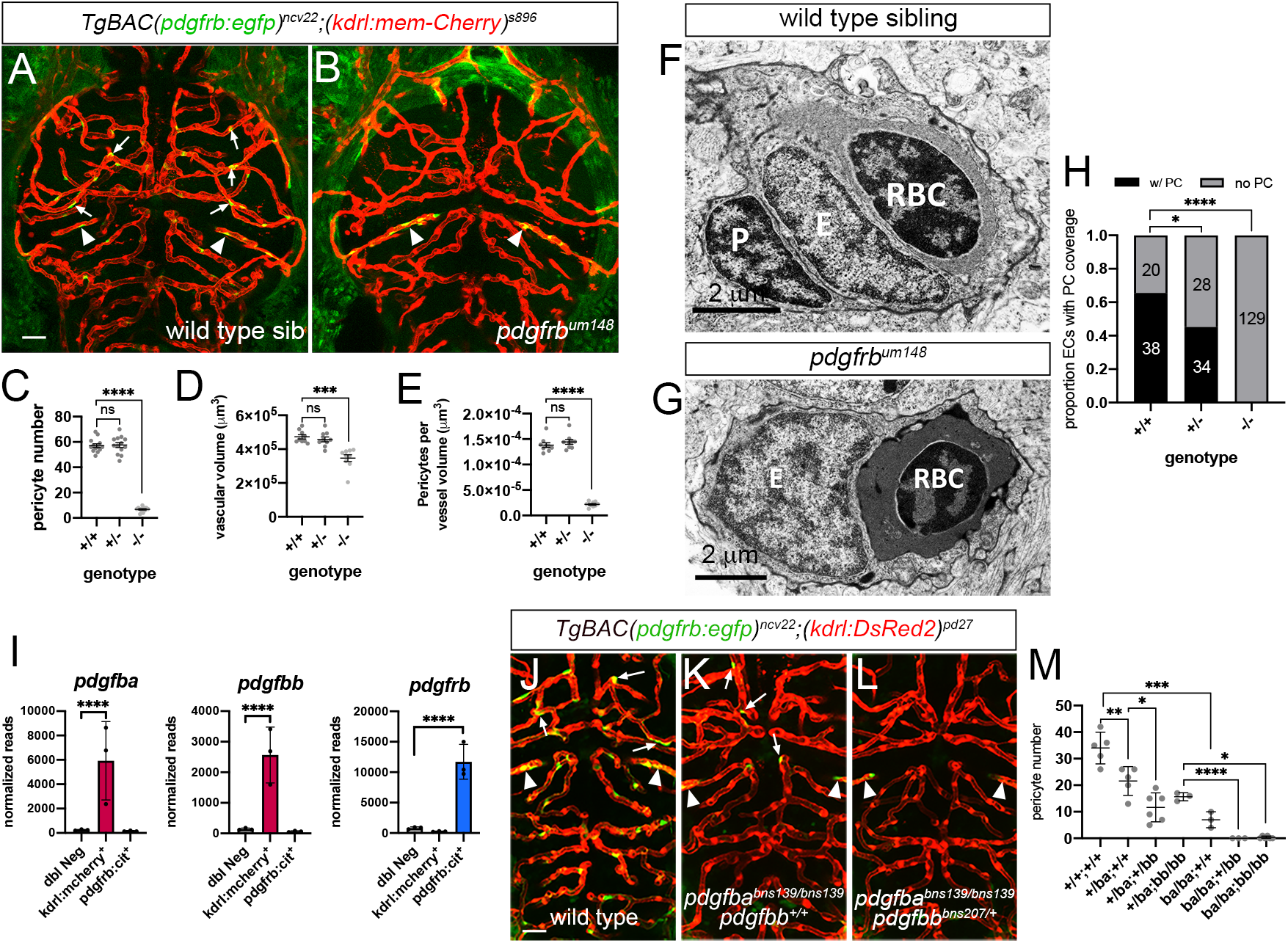
Pdgfb signaling is essential for brain pericyte development. (**A, B**) Confocal micrographs of central arteries in (**A**) wild type or (**B**) *pdgfrb^um148^* mutants bearing *TgBAC(pdgfrb:egfp)^ncv22^* (green) and *Tg(kdrl:memCherry)^s896^* (red) transgenes at 5 dpf. Arrows denote selected brain pericytes; metencephalic artery indicated by arrowheads. (**C**) Pericyte numbers in embryos of indicated genotype. n=13 per genotype. (**D**) Vascular volume in indicated genotype (n=9 for each genotype), data not normally distributed, analysis of variance (P=0.0002), multiple comparisons by Kruskal-Wallis. (**E**) Pericytes per vessel volume in indicated genotype (n=9 per genotype). (**C, E**) Data normally distributed; one-way ANOVA, P<0.0001; multiple comparison by Dunnett’s. (**C-E**) Quantification at 5 dpf. (**F, G**) Transmission electron microscopy (TEM) sections of cranial blood vessels from (**F**) wild type and (**G**) *pdgfrb^um148^* mutant embryos at 5 dpf. (**F, G**) Scale bar is 2 microns; P – pericyte, E – endothelial cell, RBC – red blood cell. (**H**) Quantification of pericyte coverage on cranial vessels in TEM sections. Fisher’s exact test. (**I**) Expression of indicated gene in triplicate RNA-seq libraries from indicated cell type at 5dpf. ****adjP<0.0001; see Lawson et al (2020). (**J-L**) Confocal images of (**J**) wild type, (**K**) *pdgfba^bns139^* mutant and (**L**) *pdgfba^bns139/bns139;^pdgfb^bbns207/+^* embryos at 5 dpf expressing *TgBAC(pdgfrb:egfp)^ncv22^* (green) and *Tg(kdrl:dsRed2)^pd27^* (red). Arrows denote selected brain pericytes; metencephalic artery indicated with arrowheads. (**M**) Quantification of periyctes at 5 dpf of indicated genotype. (**C-E, H, M**) ****p<0.001, ***p<0.005, **p<0.01, *p<0.05, ns – not statistically significant. (**A, B, J-L**) Dorsal views, scale bar is 30 μm.

As noted, *pdgfrb* encodes a receptor tyrosine kinase that is activated by Pdgfb expressed by endothelial cells (reviewed in Gaengel et al., 2009). In zebrafish, there are duplicate *pdgfb* genes, referred to as *pdgfba* and *pdgfbb* (**Fig. S2A**) and both show enriched expression in RNA-seq from isolated *kdrl:mcherry*-positive cells, consistent with their recently described endothelial expression in zebrafish (**Fig. 1I**; Lawson et al., 2020; Stratman et al., 2020). By contrast, *pdgfrb* is enriched in*pdgfrb:citrine* cells from the same larvae (**Fig. 1I**). To test the requirement for *pdgfb* genes in pericyte development, we generated *pdgfba^bns139^* and *pdgfbb^bns207^* mutants (**Fig. S2B**). Similar to *pdgfrb^um148^* mutants, *pdgfba^bns139^;pdgfbb^bns207^* double mutant embryos appeared normal throughout larval development (**Fig. S2C**) Both *pdgfba^bns139^* and *pdgfbb^bns207^* mutant larvae showed reduced pericyte coverage in central arteries at 4 dpf, with a weaker defect in *pdgfbb^bns139^* mutants (**Fig. 1J-M**). Furthermore, *pdgfba^bns139^* mutants with a single wild type copy of *pdgfbb^bns207^* or double mutants (**Fig. 1M**), showed more severe pericyte loss similar to *pdgrb^um148^* mutants (see **Fig. 1C**), suggesting a cooperative role for these ligands in brain pericyte development.

### Pdgfrb is required for vascular stability at post-larval stages

Despite loss of brain pericytes, *pdgfrb* mutant larvae do not exhibit hemorrhage or other circulatory defects (Ando et al., 2016; Kok et al., 2015). To determine if there are defects at later stages, we grew embryos from respective incrosses of *pdgfrb^um148^* and *pdgfrb^sa16389^* heterozygous carriers to adulthood. By 3 months of age, we noted aberrant swimming behavior in *pdgfrb* mutant fish (**Movie S1, S2**). Mutant siblings also presented with misshapen cranial morphology and discoloration (**Fig. 2A, B**). Observation of brain morphology revealed blood accumulation in *pdgfrb* mutant adults, compared to wild type siblings at 3 months (**Fig. 2C-E**). The locations of apparent bleeds were not consistent between individuals and appeared throughout the brain (**Fig. 2E**). Closer inspection of vascular morphology using an endothelial-specific transgene *(Tg(fli1:myr-mcherry)^ncy1^)* revealed at least two mechanisms for blood accumulation. First, blood vessels in these areas exhibited dilation, which likely resulted in lower velocity flow and blood pooling (**Fig. 2E-E”, arrow indicates same region in all three panels**). Second, we observed blood in extra-vascular space, which is defined as a hemorrhage (**arrowheads in Fig. 2E-E”**). Furthermore, we observed both of these defects as early as 30 dpf in *pdgfrb* mutant fish (**Fig. S3**).

**Figure 2.**
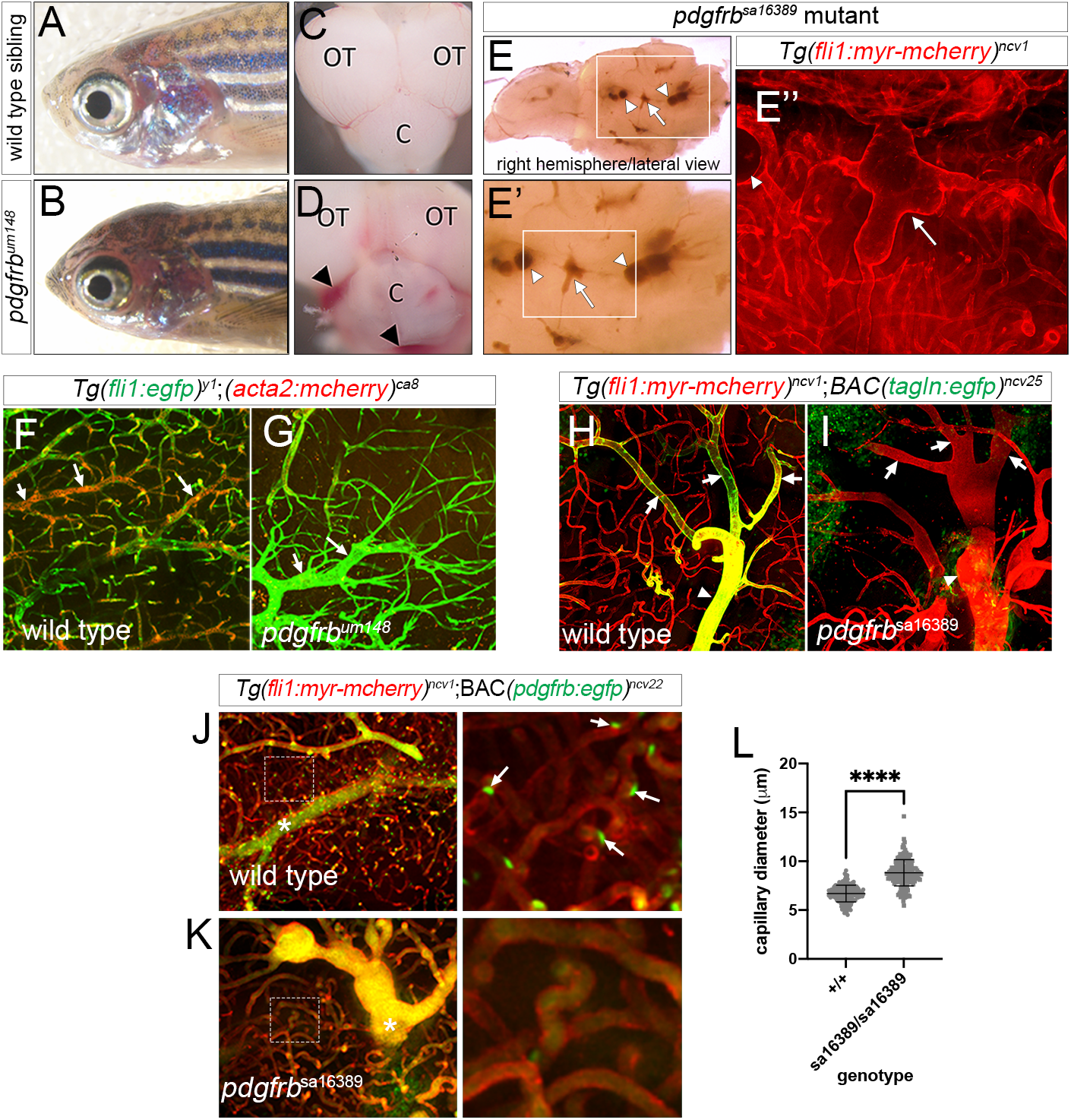
Vascular instability in *pdgfrb* mutant brains. (**A-D**) Transmitted light images of 3 month old (**A**) wild type and (**B**) *pdgfrb^um148^* mutant zebrafish heads; lateral view, anterior to right and brains dissected from (**C**) wild type and (**D**) *pdgfrb^um148^* mutant zebrafish; dorsal view, anterior is up. Arrowheads denote areas of blood accumulation. OT – optic tectum, C – cerebellum. (**E**) Lateral view of the right hemisphere from a *pdgfrb^sa16389^* mutant brain. Boxed region magnified in (**E’**). (**E’’**) Boxed region in (E’), endothleial cells visualized with *Tg(fli1:myr-mcherry)^ncv1^* transgene. (**E-E’’**) Arrow denotes blood accumulation in dilated arteriole; arrowheads denote hemorrhages. (**F, G**) Two-photon micrographs of *Tg(fli1a:egfp)^y1^* (endothelial cells, green) and *Tg(acta:mcherry)^ca8^* (vascular smooth muscle cells [VSMC], red) in (**F**) wild type and (**G**) *pdgfrb^um148^* mutant blood vessels in OT at 3 months. Arrows denote arteriole. (**H, I**) Confocal images of *TgBAC(tagln:egfp)^ncv25^* (VSMC, green) and *Tg(fli1:myr-mCherry)^ncv1^* (red) in forebrain vasculature of (**H**) wild type or (**I**) *pdgfrb^sa16389^* mutant. Arrows denote arteriole branches, arrowhead is arterial trunk in same anatomical region. Scale bars, 50 μm. (**J, K**) Confocal images of *TgBAC(pdgfrb:egfp)^ncv25^* (green) and *Tg(fli1:myr-mCherry)^ncv1^* in forebrain vasculature of (**J**) wild type or (**K**) *pdgfrb^sa16389^* mutant. Boxed areas denote magnified views to the right. Autofluorescence from circulating blood is indicated by asterisks. Scale bars, 100 μm or 20 μm (enlarged view). (**L**) Quantification of forebrain capillary diameter at 3 months in adults of indicated genotype. Error bars are mean with SD of at least 80 capillary diameter measuerments each from 4 animals. Data are not normally distributed; ****p<0.0001 by Mann-Whitney test.

Blood vessel dilation and hemorrhage may occur due to reduced VSMC coverage, which provides structural support for larger caliber arteries. Major arterioles in the optic tectum show expression of *acta2:mcherry* in a wild type adult brain at 3 months (**Fig. 2F**). By contrast, arterioles in the same location in a *pdgfrb^um148^* mutant sibling fail to express *acta2:mcherry* and appear dilated (**Fig. 2G**). We noted similar VSMC loss in *pdgfrb^sa16389^* mutants, visualized using *tagln:egfp* (**Fig. 2H, I**). For example, an arterial trunk with three branches showed extensive VSMC coverage in a wild type individual (**Fig. 2H**). By contrast, *pdgfrb^sa16389^* mutant arteries at the same anatomical location lacked *tagln:egfp-positive* cells and were dilated (**Fig. 2I**). Observation of wild type and *pdgfrb^um148^* mutant siblings as early as 45 dpf revealed loss of *acta2:mcherry* expression, with arterial dilation and hemorrhage (**Fig. S3C-H**). In addition to VSMC defects, we observed a loss of pericytes from brain capillaries compared to wild type siblings at 3 months (**Fig. 2J, K**) and an associated increase in capillary diameter (**Fig. 2L**). Together, these observations suggest a requirement of Pdgfrb for VSMCs development, concomitant with vascular stabilization between juvenile and adult stages in the zebrafish.

### Loss of Pdgfb signaling selectively affects vascular smooth muscle development during embryogenesis

Based on VSMC loss in *pdgfrb* mutant adults, we analyzed larval stages for defects in VSMC coverage. Consistent with previous observations using *Tg(acta2:mcherry)^ca8^* (Whitesell et al., 2014), we observed VSMC coverage predominantly on the ventral wall of the dorsal aorta at 5 dpf (**Fig. 3A**). However, we did not observe any difference in the number of *acta2:mcherry-positive* on the dorsal aorta in *pdgfrb^um148^* mutants (**Fig. 3B, C**). VSMC coverage of the ventral aorta at 4 dpf appeared similarly unaffected by loss of *pdgfrb* (**Fig. 3D-F**). We next assessed VSMC coverage at the CoW, where we observed *acta2:mcherry*-positive VSMC at 5 dpf in wild type siblings (**Fig. 3G**). By contrast, *pdgfrb^um148^* mutant embryos exhibit a significant decrease in CoW VSMCs (**Fig. 3H, I**). We find a similar loss of VSMC coverage in embryos lacking *pdgfba* and *pdgfbb*, as assessed using *pdgfrb:egfp*, which is co-expressed with *acta2:mcherry* at this stage in CoW VSMCs (**Fig. 3J-L; see Fig. S1C**). Thus, zebrafish VSMCs exhibit anatomically distinct requirements for signaling through Pdgfrb.

**Figure 3.**
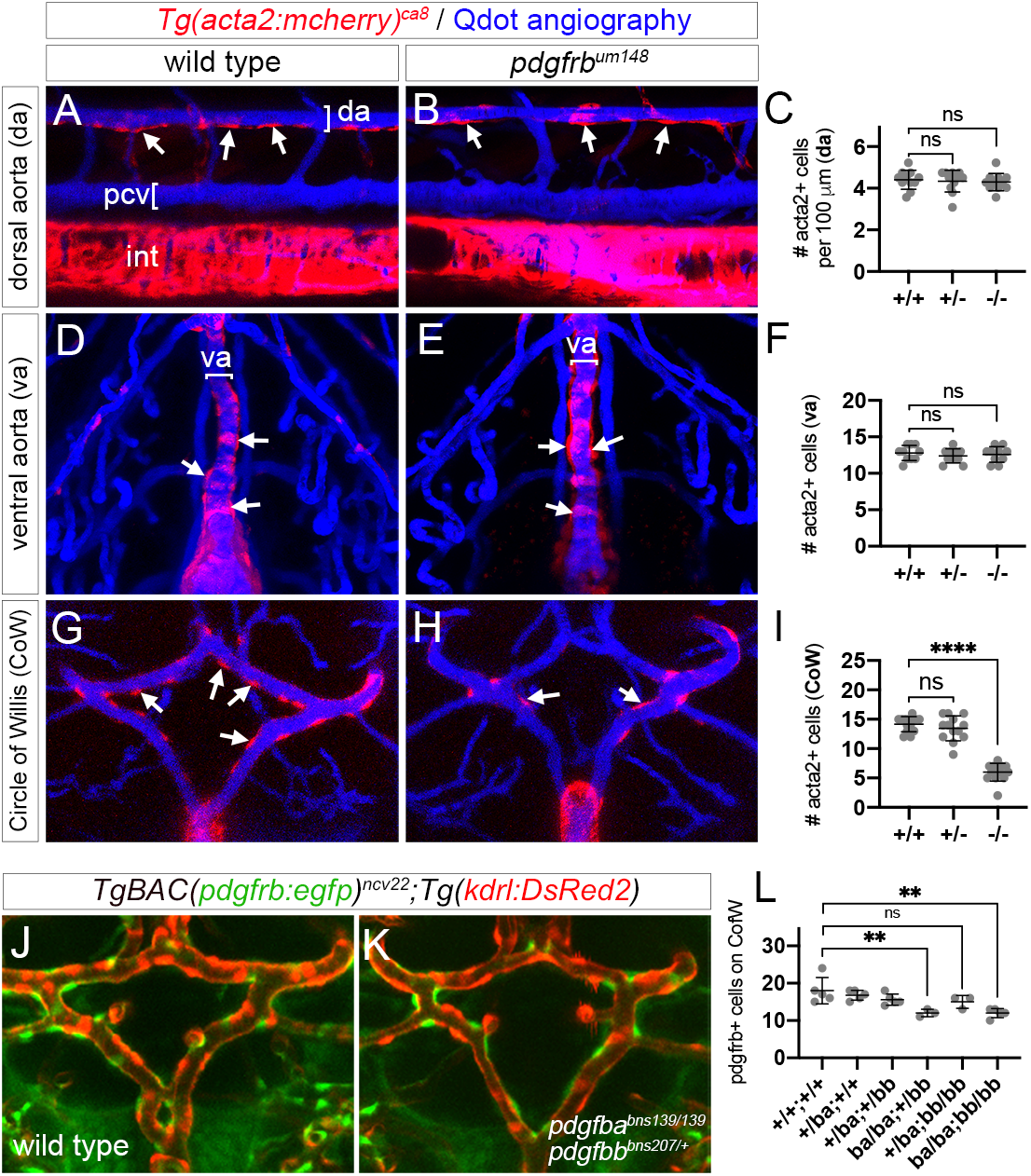
Pdgfrb is selectively required for embryonic vascular smooth muscle development. (**A, B, D, E, G, H**) Confocal images of *Tg(acta2:mcherry)^ca8^* (red, VSMCs) larvae subjected to microangiography to visualize patent blood vessels (blue). Arrows denote selected VSMCs. (**A, B**) VSMC on dorsal aorta (da) in (**A**) wild type and (**B**) *pdgfrb^um148^* mutants at 5 dpf. pcv – posterior cardinal vein, int – intestine. Lateral view, anterior to left, dorsal is up. (**D, E**) VSMC on ventral aorta (va) in (**D**) wild type and (**E**) *pdgfrb^um148^* mutants at 4 dpf. Ventral view, anterior is up. (**G, H**) VSMC on Circle of Willis in (**G**) wild type and (**H**) *pdgfrb^um148^* mutants at 5 dpf. Dorsal view, anterior is up. (**C, F, I**) Quantification of VSMCs on (**C**) da, (**F**) va, and (**I**) CoW in larvae of indicated genotype. (**C, I**) Data not normally distributed. Analysis of variance using Kruskal-Wallis test (not significant for da; p<0.0001 for CoW), multiple comparisons using Dunn’s, ****p<0.0001, ns – not statistically significant. (**F**) Data normally distributed, no significant differences by one-way ordinary ANOVA (p=0.6873). (**J, K**) Confocal images of the CoW in (**J**) wild type and (**K**) *pdgfba^bns139/bns139^;pdgfbb^bns207/+^* mutant embryos bearing *TgBAC(pdgfrb:egfp)^ncv22^; (kdrl:dsred2)^pd27^*. (**L**) Quantification of *pdgfrb:egfp-positive* cells on CoW. Data normally distributed; one-way ANOVA, p=0.0008; Tukey’s multiple comparison test, **p<0.01, ns – not statistically significant.

### Pdgfra does not play a compensatory role in trunk VSMC development

Development of trunk VSMC in *pdgfrb^um148^* mutant zebrafish is similar to Pdgfb mutant mouse embryos (Hellstrom et al., 1999). However, zebrafish expressing a dominant negative Pdgfrb show reduced VSMC coverage at the dorsal aorta (Stratman et al., 2017). Dominant negative proteins can interfere with related molecules, while nonsense mediated decay of the *pdgfrb^um148^* transcript may upregulate compensatory paralogous genes (El-Brolosy et al., 2019) during trunk VSMC development. A candidate in this regard is the related Pdgfra receptor (Andrae et al., 2008), which is enriched in *pdgfrb:citrine* positive cells (**Fig. S4**). Notably, *pdgfra* and *pdgfrb* are known to play compensatory roles during zebrafish and mouse craniofacial development (McCarthy et al., 2016). Therefore, we assessed VSMCs in embryos lacking both *pdgfra* and *pdgfrb*. From an incross of *pdgfra^b1059/+^;pdgfrb^um148/+^* carriers, we observed that approximately one-half of *pdgfra^b1059^* mutants displayed severe edema around the heart and gut, as well as the forebrain, concomitant with loss of blood circulation at 4 dpf (**Fig. 4A, E, Table S1**). Remaining mutant siblings exhibited jaw defects, as previously observed (Eberhart et al., 2008), but normal circulatory flow (**Fig. 4B, E, Table S1**). By contrast, *pdgfrb^um148^* mutant siblings, including those heterozygous for *pdgfra^b1059^* or doubly heterozygous, were normal (**Fig. 4C-E**). Notably, loss of *pdgfrb* did not increase the penetrance of circulatory defects or edema in *pdgfra* mutants (**Fig. 4E, Table S1**). We also noted focal hemorrhages, which occurred at very low penetrance in single mutants for *pdgfra^b1059^* and were typically located ventral to the eye (**Fig. 4F, G, Table S2**). The penetrance of hemorrhage increased slightly with the loss of one or two alleles of wild type *pdgfrb*, but was never observed in *pdgfrb^um148^* mutants with homozygous wild type *pdgfra* (**Fig. 4F, Table S2**).

**Figure 4.**
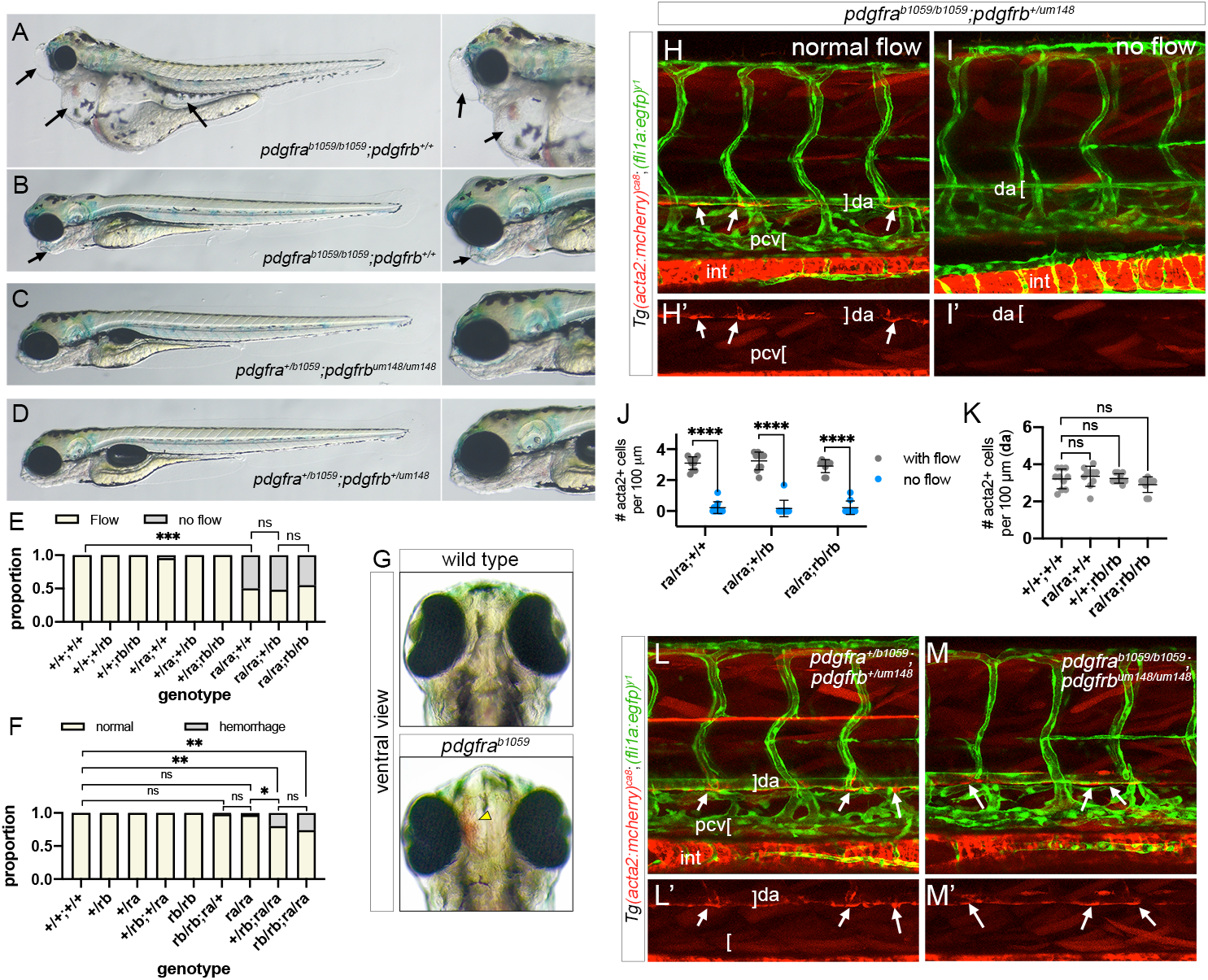
Pdgfra does not compensate for Pdgfrb deficiency during vascular smooth muscle development. (**A-D**) Transmitted light images of embryos of the following genotype at 5 dpf: (**A, B**) *pdgfra^b1059/b1059^;pdgfrb^+/+^*, (**C**) *pdgfra^+/b1059^;pdgfrb^um148/148^*, (**D**) *pdgfra^+/b1059^;pdgfrb^um148^*. Lateral views, dorsal is up, anterior to left. (**A**) Arrows denote edema. (**B**) Arrow indicates jaw. (**E**) Proportion of embryos of indicated genotype with or without blood circulation. (**F**) Proportion of embryos of indicated genotype with or without hemorrhage. (**E, F**) Fisher’s exact test, *p<0.05, **p<0.005, ***p<0.0005, ns – not significant. (**G**) Transmitted light images of wild type and *pdgfra^b1059^* mutant siblings at 5 dpf. Ventral views, anterior is up. Arrowhead indicates hemorrhage. (**H, I, L, M**) Confocal images of trunk vessels in (**H,I**) *pdgfra^b1059/b1059^;pdgfrb^+/um148^*, (**L**) *pdgfra^+/b1059^;pdgfrb^+/um148^*, and (**M**) *pdgfra^b1059/b1059^;pdgfrb^um148/um148^* larvae bearing *Tg(acta2:mcherry)^ca8^* (red, VSMC) and *Tg(fli1a:egfp)^y1^* (green, endothelial cells). Embryos in (**H, L,** and **M**) have normal circulatory flow; embryo in (**I**) has no flow. (**H’, I’, L’, M’**) Red channel showing VSMC coverage on dorasal aorta (da) for each corresponding overlay panel; arrows denote selected VSMCs. pcv – posterior cardinal vein, int – intestine. (**J, K**) Number of VSMCs per 100 μm da in embryos of indicated genotype. (**J**) Embryos with or without flow (n=10 individual embryos for each class). Paired t-test, ****p<0.00001. (**K**) Only embryos with circulation considered. Data not normally distributed. Analysis of variance by Kruskal-Wallis (P=0.1035); no statistically significant comparisons (ns).

We next assessed VSMC coverage on the dorsal aorta in *pdgfra;pdgfrb* larvae. Previous studies have shown that circulation through the trunk vasculature is essential for acquisition of dorsal aorta VSMC (Chen et al., 2017). Accordingly, we observed a loss of dorsal aorta VSMCs in *pdgfra^b1059^* mutant larvae without flow at 4 dpf, while genotypically identical siblings with circulation appeared normal (**Fig. 4H-J**). Patterning of the trunk blood vessels was otherwise relatively normal (**Fig. 4H, I**). We subsequently restricted our analyses to embryos with normal circulation. In these cases, we did not observe any significant decrease in the numbers of VSMC on the dorsal aorta of *pdgfra^b1059^;pdgfrb^um148^* double mutant embryos at 4 dpf compared to other genotypes, including wild type (**Fig. 4K-M**). These results suggest that *pdgfra* and *pdgfrb* are dispensable for initial specification and recruitment of VSMCs at the dorsal aorta at this stage.

### Pdgfrb is dispensable for mural cell coverage of large caliber trunk vessels

In *pdgfrb* mutant larvae, we observed a modest cranial VSMC defect that appears to be more severe at adult stages. Therefore, we assessed the possibility that trunk MC populations may be similarly affected at adult stages. At 3 months of age, both wild type and *pdgfrb^sa16389^* mutant individuals showed similar degrees of coverage with *pdgfrb:egfp-positive* MCs in the caudal artery (CA) and posterior cardinal vein (PCV; **Fig. 5A-C**). We observed extensive pericyte coverage on small caliber capillaries within muscle trunk tissue, as well as MC coverage on arterioles in wild type siblings (**Fig. 5D-F**). By contrast, capillaries lacked any *pdgfrb:egfp-positive* pericytes in *pdgfrb^sa16389^* mutants, although MCs persisted on arterioles and larger caliber arteries (**Fig. 5G-I, J-L**). We noted that trunk capillaries were slightly dilated (**Fig. 5M**). Otherwise, the overall vascular anatomy in the trunk region was relatively normal in *pdgfrb^sa16389^* mutants and focal distensions (microaneurysms) similar to those in brain capillaries (see **Fig. 2E**) were rarely observed.

**Figure 5.**
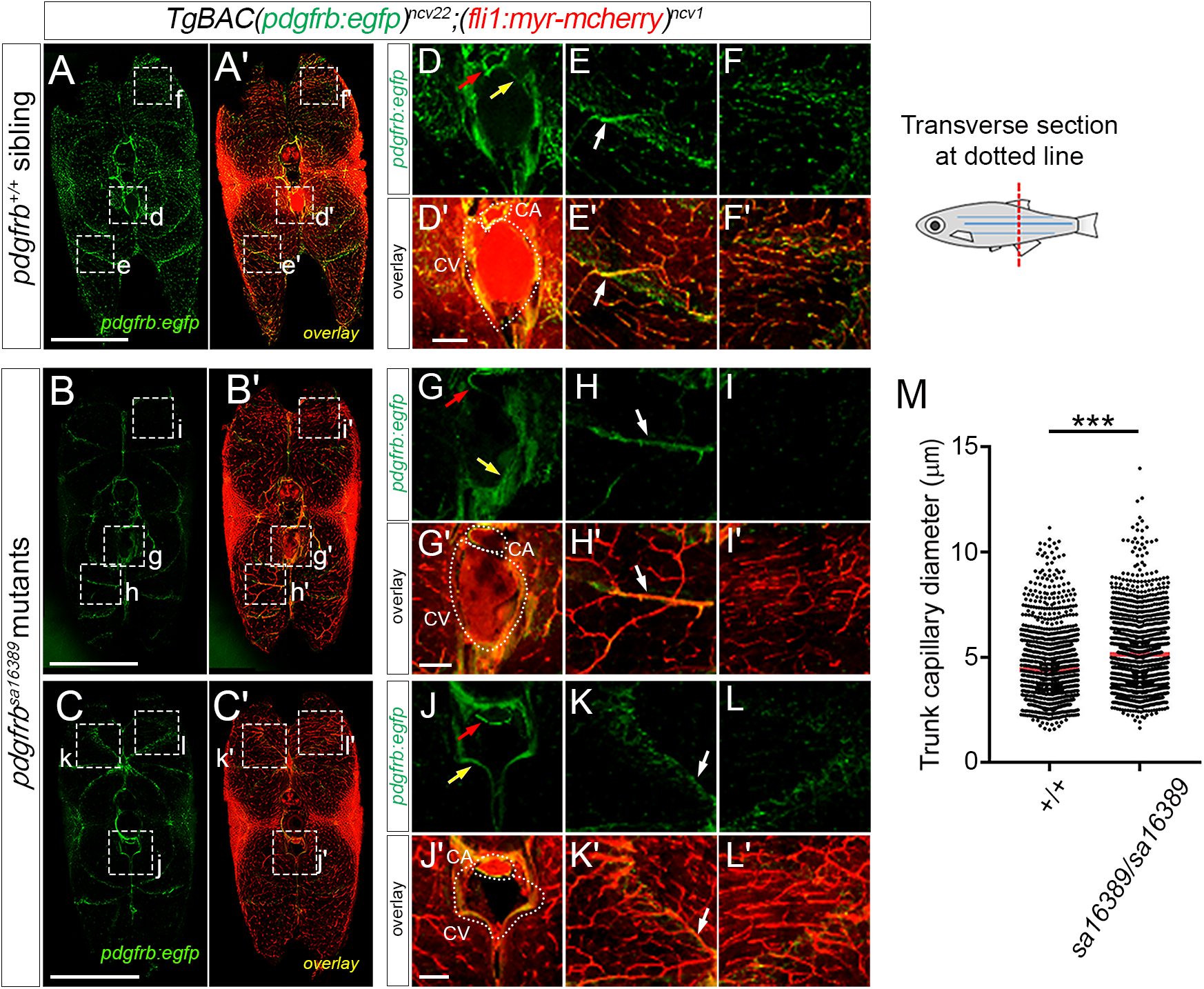
Trunk vasculature of *pdgfrb* mutants at 3 months. (**A-L**) Confocal images of trunk vessels in (**A, D-F**) wild type or (**B, C, G-L**) *pdgfrb^sa16389^* mutants bearing *TgBAC(pdgfrb:egfp)^ncv22^* (mural cells, green) and *Tg(fli1:myr-mCherry)^ncv1^* (endothelial cells, red) at 3 months from cross-section (300 μm-thick) through caudal region, as depicted at right. Boxed areas (“d-f, “g-l”) magnified to the right. Red arrows, MC on caudal artery. Yellow arrows, MCs on caudal vein. White arrows, MCs on arteriole. Scale bars, 1 mm or 100 μm (enlarged view). (**M**) Quantification of trunk capillary diameter in wild type or *pdgfrb^sa16389^* mutants at 3 months. Lines and dots indicate average and value of each capillary diameter from 4 animals, respectively. More than 80 points of capillary diameter were randomly measured in individual zebrafish. ***p<0.001, significant difference between two groups.

### The pronephric glomerulus lacks mesangial cells in *pdgfrb* mutants

In mouse, *Pdgfb* or *Pdgfrb* deficiency leads to severe defects in kidney development due to the failure to form *Pdgfrb-positive* mesangial cells (Levéen et al., 1994; Soriano, 1994). Therefore, we investigated the glomerular architecture in zebrafish *pdgfrb* mutants by TEM. In pdgfrbum148/+ heterozygous larvae, pronephric glomerular capillary endothelial cells, podocytes, and mesangial cells were readily identified at 4 dpf (**Fig. 6A-F**) using previously described criteria (Sakai and Kriz, 1987). For mesangial cells, we observed an irregular-shaped cell surrounded by extracellular matrix, cytoplasmic processes extending between the basement membrane and the fenestrated endothelium, and a prominent nucleus (**Fig. 6C, E, F**). By contrast, *pdgfrb^um148^* mutant glomeruli showed a simplified architecture with fewer cells, dilated capillaries and the absence of discernable mesangial cells (**Fig. 6G**). However, podocytes, fenestrated endothelial cells, and their intervening glomerular basement membranes were still observed (**Fig. 6G**). These structural changes are reminiscent of the glomerular phenotype in *Pdgfb* and *Pdgfrb* mutant mice suggesting a conserved role in mesangial cell development (Levéen et al., 1994; Lindahl et al., 1998; Soriano, 1994).

**Figure 6.**
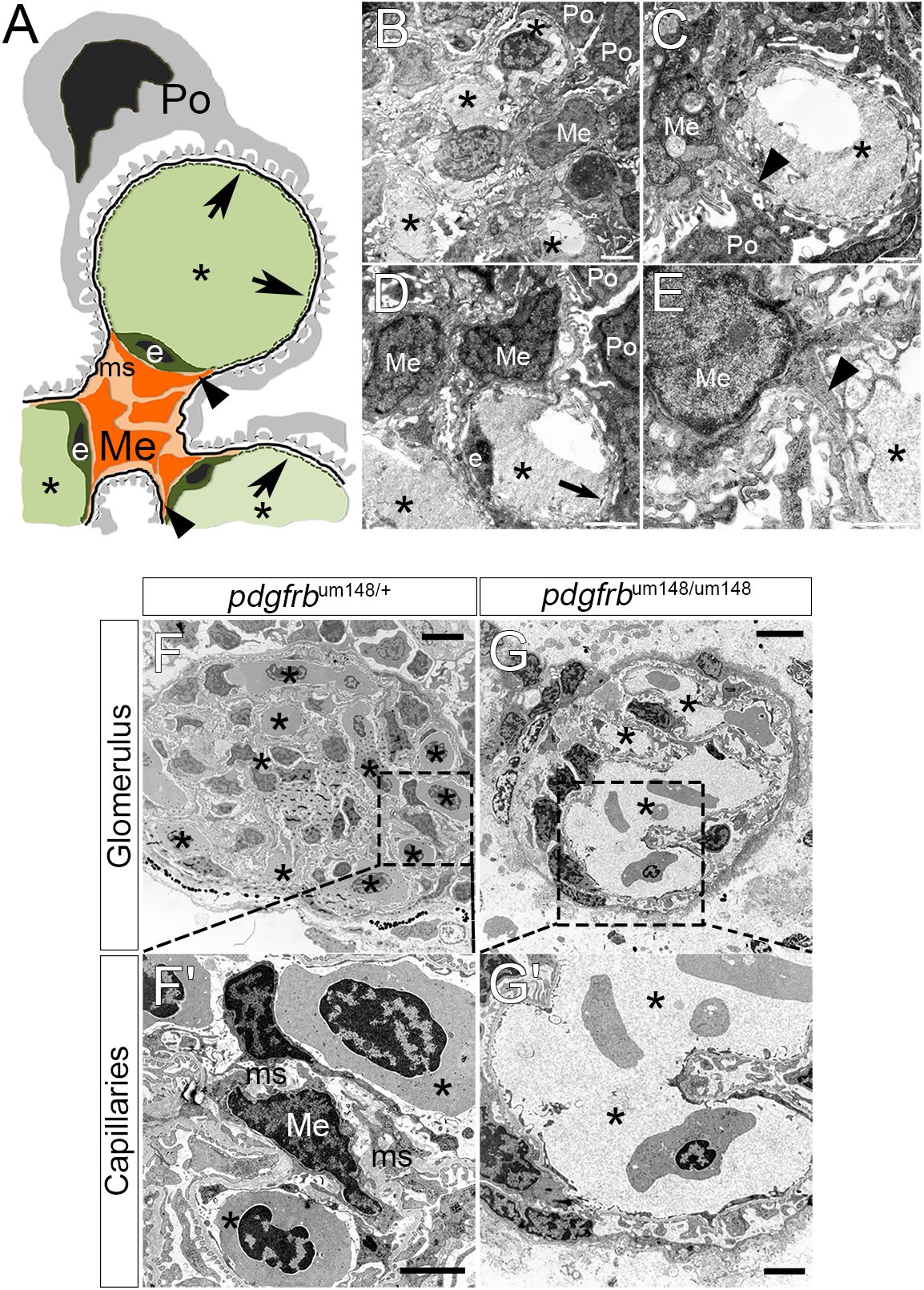
Mesangial cells in *pdgfrb* mutants. (**A**) Schematic of glomerular tuft. Fenestrated ECs (E) lining capillary lumen (*) and mesangial cells (Me) found within the mesangium (ms) shown on blood side of glomerular basement membrane (GBM; black line). Podocytes (Po) and their foot processes are on urinary side of GBM. (**B-E**) Electron micrographs of transverse sections of 4 dpf zebrafish pronephric glomerulus. Mesangial cells (Me) are identified on the blood side of the GBM. Arrowheads in **A, C**, and **E** show mesangial processes that embed between the glomerular ECs on one side and the GBM. Arrows in **A** and **D** show the fenestrated ECs of the glomerular tuft. (**F, G**) TEM micrographs of transverse sections of mesonephric glomeruli from adult (**F**) *pdgfrb^um148/+^* and (**G**) *pdgfrb^um148^* mutant fish. Ultra-structurally, mutants exhibit large aneurysmal capillaries (*) and absence of mesangial cells (Me). (**F’, G’**) Higher magnification images of areas denoted in (**F, G**).

### The coronary vasculature lacks mural cells in adult *pdgfrb* mutants

Zebrafish coronary vessels develop from 1 to 2 months of age by angiogenic sprouting of endothelial cells from the atrioventricular canal (Harrison et al., 2015). Although pericytes in coronary capillaries have been reported (Hu et al., 2001), it is unclear when MC coverage of coronary vessels begins. Already at 2 months of age, we found that all coronary vessels, including those at the angiogenic front, were covered by *pdgfrb:egfp-positive* MCs (**Fig. 7A**). This indicates that MCs are recruited to the newly formed vessels already during angiogenic expansion of coronary vessels. By contrast, *pdgfrb^sa16389^* mutants lacked coronary vessel MCs at 2 months of age (**Fig. 7B**). In addition, the angiogenic front of the coronary endothelial network was reduced in *pdgfrb^sa16389^* mutants (**Fig. 7B**). By 4 months, the coronary vessel network in *pdgfrb^sa16389^* mutants continued to be sparser than wild type siblings and capillaries still lacked MC coverage (**Fig. 7C-F**). At 8 months, wild type coronary vessels had developed further to cover the ventricle completely (**Fig. 7G**). In 8-month-old *pdgfrb^sa16389^* mutants, the coronary vasculature had further expanded compared to 4-months, but was still sparser than in wild type, and appeared immature (**Fig. 7G-I**). Moreover, *pdgfrb^sa16389^* mutants showed areas where the vasculature appeared partially disconnected and displayed abnormal capillary loops (**Fig. 7H**). These results suggest that coronary MCs are required for proper coronary vessel development in zebrafish, as previously suggested (Mellgren et al., 2008).

**Figure 7.**
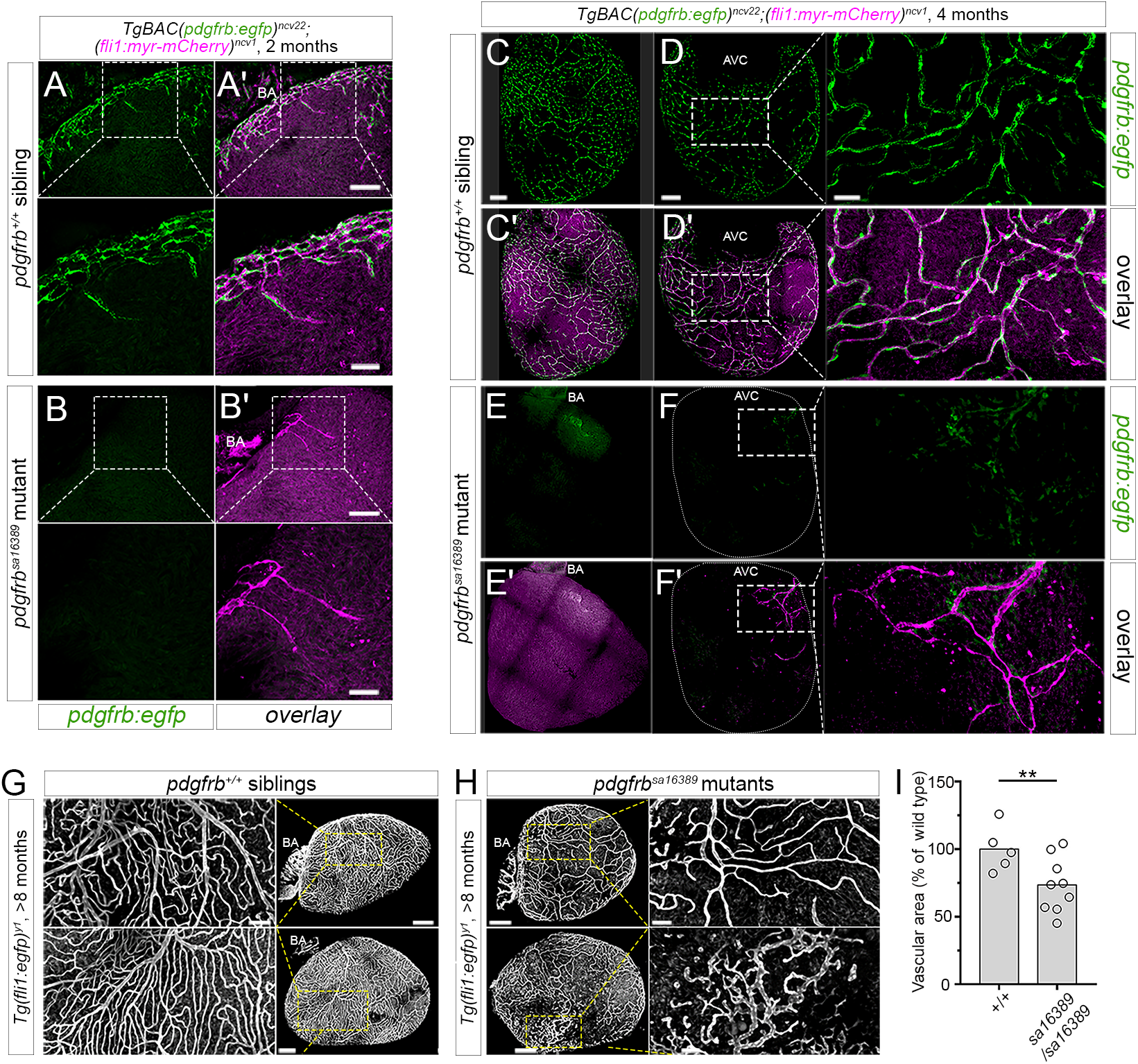
Coronary vessel defects in *pdgfrb* mutants. (**A-F**) Confocal images of coronary vessels on ventricular wall showing *TgBAC(pdgfrb:egfp)^ncv22^* expression in (**A, C, D**) wild type and (**B, E, F**) *pdgfrb^sa16389^* mutant siblings at (**A, B**) 2 months and (**C-F**) 4 months. (**A’-F’**) Overlay of images from *pdgfrb:egfp* (MCs, green) and *Tg(fli1:myr-mCherry)^ncv1^* (endothelial cells, magenta). Scale bars are (**A, B**) 100 μm, (**C, E**) 40 μm, or (D, F) 20 μm. Coronary vessels on ventricular wall facing (**C, E**) atrium or (**D, F**) opposite wall. White dotted lines in *pdgfrb^sa16389^* mutant depict ventricle shape. Boxes indicate magnified areas. AVC, atrioventricular canal. BA, bulbus arteriosus. Scale bars, 100 μm (left and center) or 20 μm (enlarged view). (**G, H**) Confocal images of coronary vessel endothelial cells on ventricular wall facing pericardial cavity in **(G)** wild type or **(H)** *pdgfrb^sa16389^* mutants with *Tg(fli1a:egfp)^y1^* at 8 months. Boxed areas are magnified to left or right of original images. Scale bars, 200 μm or 50 μm (enlarged view). Bars and circles indicate average and value of each vascular area in ventricular wall facing pericardial cavity, respectively (right). **p<0.01.

### Liver sinusoids show normal stellate cell coverage in adult *pdgfrb* mutants

Hepatic stellate cells are viewed as the pericytes of the liver sinusoid. In contrast to other MCs, Pdgfra is constitutively expressed in quiescent hepatic stellate cells, while Pdgfrb is increased in activated hepatic stellate cells, which are regarded as the major Pdgfrb-positive cell type in the liver (Chen et al., 2008). In mice, Pdgfrb signaling is dispensable for stellate cell recruitment to sinusoids, as demonstrated in both *Pdgfb* and *Pdgfrb* null mutants (Hellstrom et al., 1999). In zebrafish, we detected *pdgfrb*:egfp-positive cells in direct contact with sinusoidal endothelial cells in the liver, suggesting that these are likely hepatic stellate cells. Furthermore, these cells were not reduced in number in *pdgfrb^sa16389^* mutants (**Fig. 8A,B**), suggesting that Pdgfrb signaling is dispensable for hepatic stellate cell recruitment in zebrafish, similar to mouse (Hellstrom et al., 1999).

**Figure 8.**
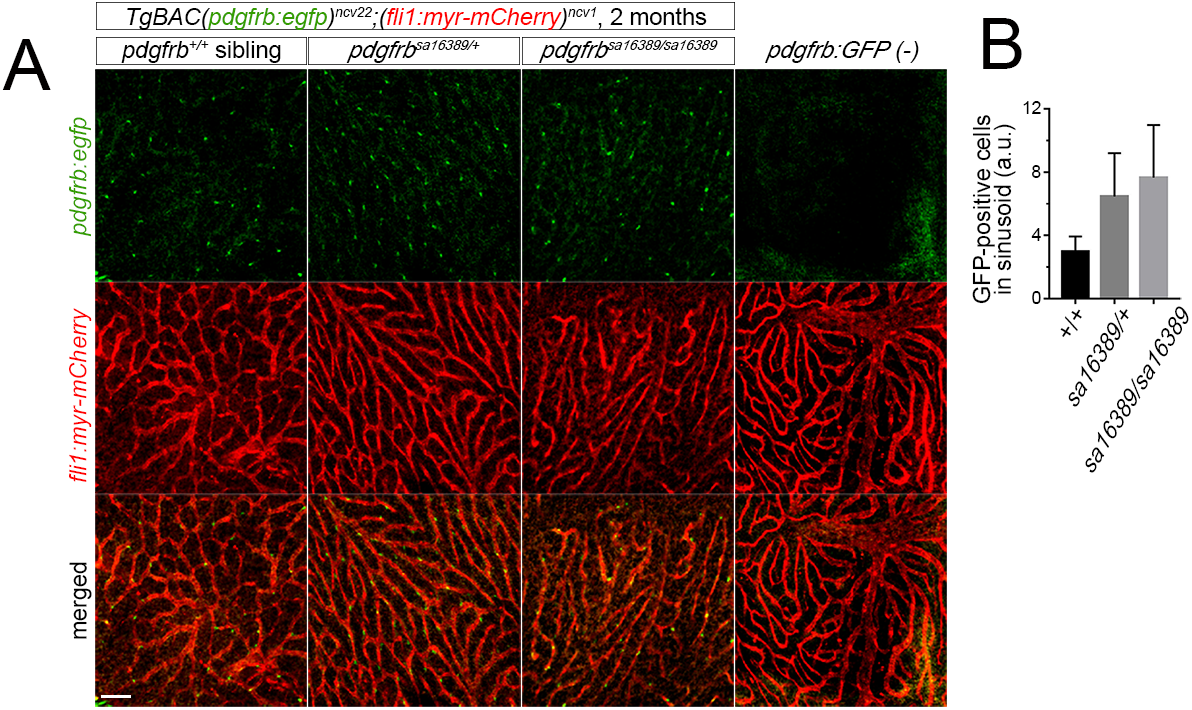
Hepatic stellate cells in *pdgfrb* mutants. (**A**) Confocal images of liver sinusoidal endothelial cells in wild type, heterozygous or homozygous *pdgfrb^sa16389^* mutant with *TgBAC(pdgfrb:egfp)^ncy22^;Tg(fli1:myr-mCherry)^ncv1^* background at 2 months. Most right column shows sinusoid of heterozygote with *Tg(fli1:myr-mCherry)^ncv1^* but without *TgBAC(pdgfrb:egfp)^ncv22^* background. Scale bar, 40 μm. (**B**) Quantification of *pdgfrb*:egfp-positive cell number divided by the volume of 3D images of the randomly observed sinusoid. The graph shows mean ± s.e.m. (n≥ 3).

## DISCUSSION

In this study, we used genetic approaches to assess the role of Pdgfb-Pdgfrb signaling for MC development and blood vessel maturation in zebrafish. By analyzing different tissues at multiple stages, we show that Pdgfb-Pdgfrb signaling is required for MC recruitment to blood vessels in the brain, trunk, glomerulus, and heart, but not for the recruitment of hepatic stellate cells to the liver sinusoids. In marked contrast to mice, zebrafish *pdgfb* and *pdgfrb* null mutants reach adulthood, in spite of extensive loss of MCs and resulting cerebral hemorrhage and edema. This difference provides a unique opportunity to better study early MC requirements and endothelial crosstalk without the confounding effects of hemorrhage and hypoxia.

The organotypic pattern of MC defects in *pdgfrb* mutant zebrafish largely parallels what has been reported for *Pdgfrb*/*Pdgfb* null mice. In the zebrafish brain parenchyma, all MC recruitment appears to be accomplished through migration and subsequent proliferation of MCs that emerged *de novo* at the cerebral base vasculature, or around the choroidal vascular plexus. The MC development in the CA and intersegmental vessels, i.e. the major arteries or arterioles covered by VSMC in the zebrafish trunk, occurred independently of Pdgfb-Pdgfrb signaling. These vessels acquire their original MC coverage through *de novo* differentiation of surrounding naïve mesenchymal cells (Ando et al., 2016; Ando et al., 2019). Therefore, the differences in the degree of MCs loss between brain and trunk vasculature of *pdgfrb* mutant zebrafish fits the proposed function of Pdgfrb signaling, namely an indispensable role in MC expansion, but not for the primary induction of MCs. The glomerular capillary phenotype in *pdgfrb* mutants also phenocopies that observed in *Pdgfb* and *Pdgfrb* null mice, suggesting an evolutionarily conserved function of Pdgfb-Pdgfrb signaling in mesangial cell recruitment. Mesangial cells are thought to participate in intussusceptive splitting of capillary loops during the formation of the glomerular tuft. The simplified glomerular structure in *pdgfrb* mutants is consistent with such a function. An alternative possibility is that mesangial cell loss leads to glomerular capillary distention due to altered hemodynamics in the absence of mesangial support. The coronary vascular phenotype in *pdgfrb* mutant zebrafish also strengthens the previous finding in mouse study (Mellgren et al., 2008) that Pdgfrb-dependent recruitment of MCs is essential for coronary vessel development. Our observation indicates that the poor coronary vascularization in the absence of MC coverage may primarily arise from defective sprouting angiogenesis, which contrasts to the brain where the reduction in vascular area may arise from destabilization/regression of established vessels. The reason why coronary angiogenesis is severely affected by MC loss is so far unclear. Finally, the hepatic stellate cell population was unaffected by the loss of *pdgfrb*, mirroring the normal appearance of these cells in *Pdgfb* or *Pdgfrb* null mice (Hellstrom et al., 1999). Thus, the importance of the Pdgfb-Pdgfrb signaling axis for MC recruitment in different organs and vascular beds appears to be conserved between mouse and zebrafish.

In spite of the similarities regarding mouse and zebrafish MC phenotypes in the absence of Pdgfrb, a major difference is the extent to which MC loss is tolerated. *Pdgfb* or *Pdgfrb* null mice die at late gestation or at birth (Levéen et al., 1994; Soriano, 1994), whereas *pdgfrb* null zebrafish survive into adulthood. In mice, microaneurysms and brain hemorrhage are observed from E11.5 onwards (Hellström et al., 2001), while similar defects occur much later in zebrafish. A likely reason for this discrepancy is differences in blood pressure. In mice, systolic left ventricular blood pressure is 2 mmHg at E9.5, but in larval zebrafish it is only 0.47 mmHg (Hu et al., 2000; Hu et al., 2001; Le et al., 2012). Zebrafish ventricular systolic blood pressure subsequently increases to 2.49 mmHg in the adult, which is comparable to that in mouse embryos (Hu et al., 2000). Hence, lower blood pressure in zebrafish larvae may potentially protect against hemorrhage despite absence of MCs. Alternatively, the different phenotypes due to MC deficiency in mice and zebrafish may relate to how oxygen is supplied to the respective embryos. In this case, pericyte deficiency may lead to hypoxia and increased production of pro-angiogenic factors, such as VEGF-A, that promote subsequent vascular abnormality and leakage (Hellström et al., 2001). Since oxygen can be taken up directly through direct gas exchange in the small zebrafish larvae, zebrafish do not typically exhibit hypoxia due to compromised circulatory function until much later stages (Kimmel et al., 1995; Rombough, 2002). In either case, the late onset of secondary effects due to MC loss in *pdgfrb* mutant zebrafish will permit more straightforward molecular analysis at embryonic stages (e.g. to investigate pericyte/endothelial crosstalk) than what is currently available in mouse.

Despite the similarities between zebrafish and mouse *pdgfrb* mutants, we noted a discrepancy with previous zebrafish studies regarding trunk VSMC development. As in mouse, *pdgfrb^um148^* display normal development of dorsal aorta VSMCs. However, previous studies over-expressing a dominant negative form of Pdgfrb in zebrafish noted decreased VSMC coverage on the dorsal aorta (Stratman et al., 2017). Since the *um148* allele causes nonsense mediated decay (Kok et al., 2015), it is possible that lack of a VSMC defect is due to genetic compensation (El-Brolosy et al., 2019). However, brain pericyte loss in *pdgfrb^um148^* mutants is fully penetrant and highly expressive. While it remains possible that there is tissue-specific compensation, it may be more likely that the dominant negative Pdgfrb used in previous studies interfered with related receptors or downstream signaling molecules to block VSMC differentiation. Alternatively, these discrepancies may reflect differences in the responsiveness of the *tagln:egfp* and *acta2:mcherry* reporter transgenes employed in the previous study and ours, respectively, to Pdgfb/Pdgfrb signaling rather than an actual loss of cells. We would note that recent studies on *pdgfba;pdgfbb* double mutants shows a similar reduction of VSMC at the dorsal aorta using the *tagln:egfp* transgene as a reporter (Stratman et al., 2020). Whether this was associated with a concomitant loss of VSMCs at the dorsal aorta using other markers or electron microscopy was not investigated. Thus, further studies are required to more definitively investigate the reasons for these differences.

Pericytes have received considerable attention in relation to the vascular abnormalities observed in several neurovascular disorders such as diabetic retinopathy, small vessel disease, and stroke (Lendahl et al., 2019). Moreover, pericyte dysfunction has been highlighted as a putative pathogenic driver in neurodegenerative diseases and aging-related cognitive decline (Sweeney et al., 2018). Mouse models of pericyte deficiency caused by genetic impairment of Pdgfb-Pdgfrb signaling have also been shown to have dysfunctional blood-brain barrier (Armulik et al., 2010; Daneman et al., 2010; Mae et al., 2021) and be a model of the rare human disease primary familial brain calcification (Arts et al., 2015; Keller et al., 2013; Nahar et al., 2019; Nicolas et al., 2013; Sanchez-Contreras et al., 2014; Vanlandewijck et al., 2015). The high degree of conservation of the mechanisms of Pdgfb/Pdgfrb-mediated MC recruitment in zebrafish may suggest that it can now be explored as a model for several of these conditions. Taking advantage of the tractability of zebrafish for chemical/genetic screening and its resistance to early death in the absence of pericytes, *pdgfrb* mutant zebrafish may prove useful in drug discovery for neurovascular diseases.

## MATERIAL AND METHODS

### Zebrafish husbandry

Zebrafish (Danio rerio) were maintained as previously described (Fukuhara et al., 2014). Embryos and larvae were staged by hpf at 28-28.5 °C. All animal experiments were performed in accordance with institutional and national regulations.

### Transgenic and mutant fish lines

Transgenic and mutant zebrafish lines were established or provided as described below. *TgBAC(pdgfrb:egfp)^ncv22^, TgBAC(pdgfrb:citrine)^s1010^, TgBAC(tagln:egfp)^ncv25^, Tg(fli1:Myr-mCherry)^ncv1^, Tg(fli1a:egfp)^yl^, Tg(kdrl:egfp)^la116^, pdgfrb^um148^* mutant, and *pdgfra^b1059^* mutant zebrafish lines were described previously (Ando et al., 2016; Choi et al., 2007; Eberhart et al., 2008; Fukuhara et al., 2014; Kok et al., 2015; Lawson and Weinstein, 2002; Vanhollebeke et al., 2015). *pdgfrb^sa16389^* mutant zebrafish were obtained from European Zebrafish Resource Center (Ando et al., 2016). *pdgfba*^bns139^ and *pdgfbb*^bns207^ mutants were generated by CRISPR/Cas9-mediated genome editing (See also **Fig. S2**). The sgRNA 5’-ggAAGGCCATAACATAAAGT-3’ was used to target *pdgfba* (ENSDARG00000086778.3), and we identified an allele carrying a 10 bp frameshift indel in the 4th exon, which encodes the conserved PDGF/VEGF homology domain. The sgRNA 5’-ggACTGCGCGGCAGACGGTTGC-3’was used to target the 3rd exon of *pdgfbb* (ENSDARG00000038139.7), and we identified an allele carrying a 26 bp frame-shift mutation and splice site deletion upstream of the region encoding the PDGF/VEGF homology domain. Guides were designed using Chopchop (Labun et al., 2019) and produced as described in Gagnon et al. (Gagnon et al., 2014) using the T7-promoter and MEGAShortscript™ Transcription kit (Invitrogen). Genotyping was performed by high resolution melt analysis using the primer pairs BA_6_fwd1: 5’-TTACAGCAGCCTGAACAGCG-3’ and BA_6_rev1: 5’-ACCCGTGCGATGTTTGATAGA-3’ for *pdgfba* and BB_3_fwd1: 5’-AGCCATCATGACAATGACTCC-3’ and BB_3_rev1: 5’-TGAGAGAATAAAAGAGAAGTGAACTGA-3’ for *pdgfbb*. For *pdgfrb^um148^*, genotyping was performed using KASP primer pairs (Biosearch Technologies) targeting the following sequence: 5’-CTGCTCTGTCTGGGCACTTCAGGTCTGGAGCTCAGTCCCAGCGCTCCACA[GATC/-]ATCCTGTCCATCAACTCGTCCTCCAGCATCACCTGCTCCGGCTGGAGTAA-3’. Genotyping for *pdgfra^b1059^* was performed as described elsewhere (Eberhart et al., 2008).

### Image acquisition by confocal microscopy and processing

Larvae were anesthetized and mounted in 1% low-melting agarose on a 35-mm-diameter glass-base dish (Asahi Techno Glass or Thermo Scientific Nunc), as previously described (Fukuhara et al., 2014). Confocal images were obtained using a FluoView FV1200 confocal upright microscope (Olympus) equipped with a water-immersion 20x (XLUMPlanFL, 1.0 NA) lens, a Leica TCS SP8 confocal microscope (Leica Microsystems) equipped with a water-immersion 25x (HCX IRAPOL, 0.95 NA) or a dry 10x (HC PLAPO CS, 0.40 NA) or a Zeiss NLO710 equipped with a 20x (W Plan-APOCHROMAT 20×/1.0, DIC D=0.17 M27 70 mm) lens. The 473 nm (for GFP), 559 nm (for mCherry), and 633 nm (for Qdot 655) laser lines in FluoView FV1200 confocal microscope and the 488 nm (for GFP) and 587 nm (for mCherry) in Leica TCS SP8 confocal microscope were employed, and 488 nm and 651 nm on the Zeiss NLO710, respectively. Where indicated, adult brain vasculature was imaged by two-photon imaging using a Zeiss NLO710 equipped with a Chameleon Ti:Sapphire pulsed laser switched between 900 and 1040 nm excitation by section to capture green and red fluorescence, respectively. Confocal or 2-photon image stacks were processed using Olympus Fluoview (FV10-ASW), Leica Application Suite 3.2.1.9702, or IMARIS 8 software (Bitplane). All images are presented as maximum intensity projections (Leica Application Suite 3.2.1.9702). Bright field images were taken with a fluorescence stereozoom microscope (SZX12, Olympus) or MZ125 microscope (Leica).

### Image acquisition by transmission electron microscopy and processing

Zebrafish embryos or adults were euthanized prior to processing. The mesonephros was dissected while the euthanized fish were on ice. The zebrafish embryos or dissected mesonephros were fixed in 2% glutaraldehyde/0.5% paraformaldehyde/0.1M cacodylate/0.1M sucrose/3 mM Cacl2 and washed in 0.1M cacodylate buffer pH 7.4 prior to staining in 2% OsO4 for 1 hour at room temperature. Samples were dehydrated and en bloc staining was performed in 2% uranyl acetate in absolute ethanol for 1 hour at room temperature. Tissue was then taken through an Epon 812/acetone series and embedded at 60°C in pure Epon 812. Thin sections of 70 nm thickness were made on a Leica EM UC6 ultramicrotome and mounted on formvar coated copper slot grids. Post-staining was done with 5% uranyl acetate pH3.5 and Venable and Cogglesall’s lead citrate. Grids were washed extensively in water. Samples were analyzed on a JEOL 1230 electron microscope.

### Statistical analysis

Data are expressed as means ± s.e.m. Statistical significance was determined by a Student’s t test for paired samples, one-way analysis of variance with Turkey’s test for multiple comparisons, or Dunnett’s Multiple Comparison Test. Data were considered statistically significant if P-values < 0.05.

## Acknowledgements

We thank to S. Schulte-Merker for providing plasmids for BAC recombineering, K. Kawakami for the Tol2 system. We also thank Sarah Childs for providing *Tg(acta2:mcherry)^ca8^* fish. We are grateful to Y. Ando, E. Nakamura, and T. Miyazaki for technical assistance. We thank Patrick White and John Polli for excellent fish care and maintenance.

## Author Contributions

K.A., C.B., and N.L. conceived and designed the research. K.A., L.E., N.L., and C.B. wrote the manuscript with significant input from all co-authors. A. G. analyzed adult brain phenotype in *pdgfrb^um148^* mutant; Y.-H. S., A. G., and D. P. analyzed phenotypes at larval stages in *pdgfrb* and *pdgfra/b* double mutants. K.A. analyzed brain, trunk, and liver vascular phenotype in *pdgfrb^sa16389^* mutants with assistance from A.C. L.E. analyzed kidney vascular phenotype. K.A. and A.C. analyzed coronary vascular phenotype. C.G., K.M., and D.S. established the *pdgfb* mutants. All authors reviewed the manuscript.

## References

Ando, K., Fukuhara, S., Izumi, N., Nakajima, H., Fukui, H., Kelsh, R. N. and Mochizuki, N. (2016). Clarification of mural cell coverage of vascular endothelial cells by live imaging of zebrafish. Development 143, 1328–1339.

Ando, K., Wang, W., Peng, D., Chiba, A., Lagendijk, A. K., Barske, L., Crump, J. G., Stainier, D. Y. R., Lendahl, U., Koltowska, K., et al. (2019). Peri-arterial specification of vascular mural cells from naïve mesenchyme requires Notch signaling. Development 146, dev165589.

Andrae, J., Gallini, R. and Betsholtz, C. (2008). Role of platelet-derived growth factors in physiology and medicine. Genes Dev 22, 1276–1312.

Armulik, A., Abramsson, A. and Betsholtz, C. (2005). Endothelial/pericyte interactions. Circulation research 97, 512–523.

Armulik, A., Genové, G. and Betsholtz, C. (2011). Pericytes: developmental, physiological, and pathological perspectives, problems, and promises. Developmental cell 21, 193–215.

Armulik, A., Genové, G., Mäe, M., Nisancioglu, M. H., Wallgard, E., Niaudet, C., He, L., Norlin, J., Lindblom, P. and Strittmatter, K. (2010). Pericytes regulate the blood–brain barrier. Nature 468, 557.

Arts, F. A., Velghe, A. I., Stevens, M., Renauld, J. C., Essaghir, A. and Demoulin, J. B. (2015). Idiopathic basal ganglia calcification-associated PDGFRB mutations impair the receptor signalling. J Cell Mol Med 19, 239–248.

Augustin, H. G. and Koh, G. Y. (2017). Organotypic vasculature: From descriptive heterogeneity to functional pathophysiology. Science 357, eaal2379.

Beck, L. and D’Amore, P. A. (1997). Vascular development: cellular and molecular regulation. The FASEB Journal 11, 365–373.

Benjamin, L. E., Hemo, I. and Keshet, E. (1998). A plasticity window for blood vessel remodelling is defined by pericyte coverage of the preformed endothelial network and is regulated by PDGF-B and VEGF. Development 125, 1591–1598.

Chen, S. W., Chen, Y. X., Zhang, X. R., Qian, H., Chen, W. Z. and Xie, W. F. (2008). Targeted inhibition of platelet-derived growth factor receptor-beta subunit in hepatic stellate cells ameliorates hepatic fibrosis in rats. Gene Ther 15, 1424–1435.

Chen, X., Gays, D., Milia, C. and Santoro, M. M. (2017). Cilia Control Vascular Mural Cell Recruitment in Vertebrates. Cell Rep 18, 1033–1047.

Choi, J., Dong, L., Ahn, J., Dao, D., Hammerschmidt, M. and Chen, J. N. (2007). FoxH1 negatively modulates flk1 gene expression and vascular formation in zebrafish. Dev Biol 304, 735–744.

Daneman, R., Zhou, L., Kebede, A. A. and Barres, B. A. (2010). Pericytes are required for blood–brain barrier integrity during embryogenesis. Nature 468, 562–566.

Eberhart, J. K., He, X., Swartz, M. E., Yan, Y. L., Song, H., Boling, T. C., Kunerth, A. K., Walker, M. B., Kimmel, C. B. and Postlethwait, J. H. (2008). MicroRNA Mirn140 modulates Pdgf signaling during palatogenesis. Nat Genet 40, 290–298.

El-Brolosy, M. A., Kontarakis, Z., Rossi, A., Kuenne, C., Gunther, S., Fukuda, N., Kikhi, K., Boezio, G. L. M., Takacs, C. M., Lai, S. L., et al. (2019). Genetic compensation triggered by mutant mRNA degradation. Nature 568, 193–197.

Farquhar, M. G. and Palade, G. E. (1962). FUNCTIONAL EVIDENCE FOR THE EXISTENCE OF A THIRD CELL TYPE IN THE RENAL GLOMERULUS: Phagocytosis of Filtration Residues by a Distinctive “Third” Cell. J Cell Biol 13, 55–87.

Fukuhara, S., Zhang, J., Yuge, S., Ando, K., Wakayama, Y., Sakaue-Sawano, A., Miyawaki, A. and Mochizuki, N. (2014). Visualizing the cell-cycle progression of endothelial cells in zebrafish. Developmental biology 393, 10–23.

Gaengel, K., Genove, G., Armulik, A. and Betsholtz, C. (2009). Endothelial-mural cell signaling in vascular development and angiogenesis. Arterioscler Thromb Vasc Biol 29, 630–638.

Gagnon, J. A., Valen, E., Thyme, S. B., Huang, P., Akhmetova, L., Pauli, A., Montague, T. G., Zimmerman, S., Richter, C. and Schier, A. F. (2014). Efficient mutagenesis by Cas9 protein-mediated oligonucleotide insertion and large-scale assessment of single-guide RNAs. PLoS One 9, e98186.

Harrison, M. R., Bussmann, J., Huang, Y., Zhao, L., Osorio, A., Burns, C. G., Burns, C. E., Sucov, H. M., Siekmann, A. F. and Lien, C.-L. (2015). Chemokine-guided angiogenesis directs coronary vasculature formation in zebrafish. Developmental cell 33, 442–454.

He, B., Chen, P., Zambrano, S., Dabaghie, D., Hu, Y., Möller-Hackbarth, K., Unnersjö-Jess, D., Korkut, G., Charrin, E., Jeansson, M., et al. (2021). Single-cell RNA sequencing reveals the mesangial identity and species diversity of glomerular cell transcriptomes Nature Communications in press.

Hellström, M., Gerhardt, H., Kalén, M., Li, X., Eriksson, U., Wolburg, H. and Betsholtz, C. (2001). Lack of pericytes leads to endothelial hyperplasia and abnormal vascular morphogenesis. The Journal of cell biology 153, 543–554.

Hellstrom, M., Lindahl, P., Abramsson, A. and Betsholtz, C. (1999). Role of PDGF-B and PDGFR-beta in recruitment of vascular smooth muscle cells and pericytes during embryonic blood vessel formation in the mouse. Development 126, 3047–3055.

Hu, N., Sedmera, D., Yost, H. J. and Clark, E. B. (2000). Structure and function of the developing zebrafish heart. The Anatomical Record 260, 148–157.

Hu, N., Yost, H. J. and Clark, E. B. (2001). Cardiac morphology and blood pressure in the adult zebrafish. The Anatomical Record 264, 1–12.

Hungerford, J. E., Owens, G. K., Argraves, W. S. and Little, C. D. (1996). Development of the aortic vessel wall as defined by vascular smooth muscle and extracellular matrix markers. Developmental biology 178, 375–392.

Isogai, S., Horiguchi, M. and Weinstein, B. M. (2001). The vascular anatomy of the developing zebrafish: an atlas of embryonic and early larval development. Dev Biol 230, 278–301.

Keller, A., Westenberger, A., Sobrido, M. J., Garcia-Murias, M., Domingo, A., Sears, R. L., Lemos, R. R., Ordonez-Ugalde, A., Nicolas, G., da Cunha, J. E., et al. (2013). Mutations in the gene encoding PDGF-B cause brain calcifications in humans and mice. Nat Genet 45, 1077–1082.

Kimmel, C. B., Ballard, W. W., Kimmel, S. R., Ullmann, B. and Schilling, T. F. (1995). Stages of embryonic development of the zebrafish. Developmental dynamics 203, 253–310.

Kok, F. O., Shin, M., Ni, C. W., Gupta, A., Grosse, A. S., van Impel, A., Kirchmaier, B. C., Peterson-Maduro, J., Kourkoulis, G., Male, I., et al. (2015). Reverse genetic screening reveals poor correlation between morpholino-induced and mutant phenotypes in zebrafish. Dev Cell 32, 97–108.

Labun, K., Montague, T. G., Krause, M., Torres Cleuren, Y. N., Tjeldnes, H. and Valen, E. (2019). CHOPCHOP v3: expanding the CRISPR web toolbox beyond genome editing. Nucleic Acids Research 47, W171–W174.

Latta, H., Maunsbach, A. B. and Madden, S. C. (1960). The centrolobular region of the renal glomerulus studied by electron microscopy. J Ultrastruct Res 4, 455–472.

Lawson, N. D., Li, R., Shin, M., Grosse, A., Yukselen, O., Stone, O. A., Kucukural, A. and Zhu, L. (2020). An improved zebrafish transcriptome annotation for sensitive and comprehensive detection of cell type-specific genes. Elife 9.

Lawson, N. D. and Weinstein, B. M. (2002). In vivo imaging of embryonic vascular development using transgenic zebrafish. Developmental biology 248, 307–318.

Le, V. P., Kovacs, A. and Wagenseil, J. E. (2012). Measuring left ventricular pressure in late embryonic and neonatal mice. Journal of visualized experiments: JoVE.

Lendahl, U., Nilsson, P. and Betsholtz, C. (2019). Emerging links between cerebrovascular and neurodegenerative diseases—a special role for pericytes. EMBO reports 20.

Levéen, P., Pekny, M., Gebre-Medhin, S., Swolin, B., Larsson, E. and Betsholtz, C. (1994). Mice deficient for PDGF B show renal, cardiovascular, and hematological abnormalities. Genes & development 8, 1875–1887.

Lindahl, P., Hellstrom, M., Kalen, M., Karlsson, L., Pekny, M., Pekna, M., Soriano, P. and Betsholtz, C. (1998). Paracrine PDGF-B/PDGF-Rbeta signaling controls mesangial cell development in kidney glomeruli. Development 125, 3313–3322.

Lindahl, P., Johansson, B. R., Levéen, P. and Betsholtz, C. (1997). Pericyte loss and microaneurysm formation in PDGF-B-deficient mice. Science 277, 242–245.

Lindblom, P., Gerhardt, H., Liebner, S., Abramsson, A., Enge, M., Hellström, M., Bäckström, G., Fredriksson, S., Landegren, U. and Nyström, H. C. (2003). Endothelial PDGF-B retention is required for proper investment of pericytes in the microvessel wall. Genes & development 17, 1835–1840.

Mae, M. A., He, L., Nordling, S., Vazquez-Liebanas, E., Nahar, K., Jung, B., Li, X., Tan, B. C., Chin Foo, J., Cazenave-Gassiot, A., et al. (2021). Single-Cell Analysis of Blood-Brain Barrier Response to Pericyte Loss. Circ Res 128, e46–e62.

McCarthy, N., Liu, J. S., Richarte, A. M., Eskiocak, B., Lovely, C. B., Tallquist, M. D. and Eberhart, J. K. (2016). Pdgfra and Pdgfrb genetically interact during craniofacial development. Dev Dyn 245, 641–652.

Mellgren, A. M., Smith, C. L., Olsen, G. S., Eskiocak, B., Zhou, B., Kazi, M. N., Ruiz, F. R., Pu, W. T. and Tallquist, M. D. (2008). Platelet-derived growth factor receptor β signaling is required for efficient epicardial cell migration and development of two distinct coronary vascular smooth muscle cell populations. Circulation research 103, 1393–1401.

Muhl, L., Genové, G., Leptidis, S., Liu, J., He, L., Mocci, G., Sun, Y., Gustafsson, S., Buyandelger, B., Chivukula, I. V., et al. (2020). Single-cell analysis uncovers fibroblast heterogeneity and criteria for fibroblast and mural cell identification and discrimination. Nature Communications 11, 3953.

Nahar, K., Lebouvier, T., Andaloussi Mae, M., Konzer, A., Bergquist, J., Zarb, Y., Johansson, B., Betsholtz, C. and Vanlandewijck, M. (2019). Astrocyte-microglial association and matrix composition are common events in the natural history of primary familial brain calcification. Brain Pathol.

Nicolas, G., Pottier, C., Charbonnier, C., Guyant-Marechal, L., Le Ber, I., Pariente, J., Labauge, P., Ayrignac, X., Defebvre, L., Maltete, D., et al. (2013). Phenotypic spectrum of probable and genetically-confirmed idiopathic basal ganglia calcification. Brain 136, 3395–3407.

Rombough, P. (2002). Gills are needed for ionoregulation before they are needed for O2 uptake in developing zebrafish, Danio rerio. Journal of Experimental Biology 205, 1787–1794.

Sakai, F. and Kriz, W. (1987). The structural relationship between mesangial cells and basement membrane of the renal glomerulus. Anatomy and Embryology 176, 373–386.

Sanchez-Contreras, M., Baker, M. C., Finch, N. A., Nicholson, A., Wojtas, A., Wszolek, Z. K., Ross, O. A., Dickson, D. W. and Rademakers, R. (2014). Genetic screening and functional characterization of PDGFRB mutations associated with basal ganglia calcification of unknown etiology. Hum Mutat 35, 964–971.

Schlondorff, D. (1987). The glomerular mesangial cell: an expanding role for a specialized pericyte. The FASEB Journal 1, 272–281.

Soriano, P. (1994). Abnormal kidney development and hematological disorders in PDGF beta-receptor mutant mice. Genes Dev 8, 1888–1896.

Stratman, A. N., Burns, M. C., Farrelly, O. M., Davis, A. E., Li, W., Pham, V. N., Castranova, D., Yano, J. J., Goddard, L. M., Nguyen, O., et al. (2020). Chemokine mediated signalling within arteries promotes vascular smooth muscle cell recruitment. Commun Biol 3, 734.

Stratman, A. N., Pezoa, S. A., Farrelly, O. M., Castranova, D., Dye, L. E., 3rd, Butler, M. G., Sidik, H., Talbot, W. S. and Weinstein, B. M. (2017). Interactions between mural cells and endothelial cells stabilize the developing zebrafish dorsal aorta. Development 144, 115–127.

Sweeney, M. D., Sagare, A. P. and Zlokovic, B. V. (2018). Blood-brain barrier breakdown in Alzheimer disease and other neurodegenerative disorders. Nat Rev Neurol 14, 133–150.

Tallquist, M. D., French, W. J. and Soriano, P. (2003). Additive effects of PDGF receptor beta signaling pathways in vascular smooth muscle cell development. PLoS Biol 1, E52.

Vanhollebeke, B., Stone, O. A., Bostaille, N., Cho, C., Zhou, Y., Maquet, E., Gauquier, A., Cabochette, P., Fukuhara, S., Mochizuki, N., et al. (2015). Tip cell-specific requirement for an atypical Gpr124- and Reck-dependent Wnt/beta-catenin pathway during brain angiogenesis. Elife 4.

Vanlandewijck, M., He, L., Mäe, M. A., Andrae, J., Ando, K., Del Gaudio, F., Nahar, K., Lebouvier, T., Laviña, B. and Gouveia, L. (2018). A molecular atlas of cell types and zonation in the brain vasculature. Nature 554, 475.

Vanlandewijck, M., Lebouvier, T., Mäe, M. A., Nahar, K., Hornemann, S., Kenkel, D., Cunha, S. I., Lennartsson, J., Boss, A. and Heldin, C.-H. (2015). Functional characterization of germline mutations in PDGFB and PDGFRB in primary familial brain calcification. PloS one 10, e0143407.

Whitesell, T. R., Kennedy, R. M., Carter, A. D., Rollins, E. L., Georgijevic, S., Santoro, M. M. and Childs, S. J. (2014). An alpha-smooth muscle actin (acta2/alphasma) zebrafish transgenic line marking vascular mural cells and visceral smooth muscle cells. PLoS One 9, e90590.

Yin, C., Evason, K. J., Asahina, K. and Stainier, D. Y. R. (2013). Hepatic stellate cells in liver development, regeneration, and cancer. The Journal of Clinical Investigation 123, 1902–1910.

